# Towards Mesoscopic Human Brain Imaging Using Non-Parametric Diffusion Tensor Distribution (DTD) MRI

**DOI:** 10.64898/2026.04.12.718025

**Authors:** Kulam Najmudeen Magdoom, Joelle E. Sarlls, Peter J. Basser

## Abstract

The majority of MR-based brain imaging methods provides macroscopic information averaged over the entire imaging voxel. Yet tissue composition and microstructure are heterogeneous within the cubic millimeter-sized MRI voxel that contains numerous distinct water pools at mesoscopic, microscopic, and nanoscopic length scales. Accurately measuring their individual characteristics in live human brain has the potential to reveal hidden salient meso/micro-structural features and uncover subtle changes that may occur in development, neurological disorders, trauma, etc. Nevertheless, because of many technical and scientific challenges, there is a dearth of robust, quantitative methods to probe tissue water dynamics at these subvoxel length scales. Here we present a novel empirical spectroscopic diffusion MRI method that estimates the probability density function (pdf) of diffusion tensors, i.e., the diffusion tensor distribution (DTD) in the human brain *in-vivo*. Our method entails performing a multi-dimensional Inverse Laplace Transform (ILT) which is generally an ill-posed and ill-conditioned problem. However, we overcome these obstacles using a hierarchy of lower-dimensional marginal distributions of the DTD estimated from diffusion weighted (DW) signals obtained from single, double, and triple pulsed-field gradient (PFG) experiments. Iteratively applying this hierarchy of marginal distribution progressively shrinks the space of admissible solutions. We extensively vet this framework with simulated DWI data obtained from realistic DTD motifs that mimic different cell and tissue properties seen in the brain. We then experimentally test our approach *in vivo* in brains of healthy normal human subjects. We segment the reconstructed DTD within a voxel to identify signatures of different tissue and cell types, and cluster these DTDs to identify various water pools. We use the high dimensional spectrum to robustly remove the free water compartment that often confounds tissue microstructure. We take ensemble averages of invariants of the micro-diffusion tensors, and measure and map their distributions to visualize salient intrinsic mesoscopic features. Since DTD MRI subsumes DTI, we also compute the family of DTI-derived quantitative imaging biomarkers from the moments of the distributions of the mean diffusivity and FA derived from the DTD. Our approach has great translational potential, revealing new microstructural features not observed previously observed in *in vivo* MRI.

## 1 Introduction

Brain tissue microstructure plays a critical role in ensuring normal function while subtle changes can serve as an early marker of emerging neuropathology. However, it is challenging to study and characterize *in vivo* since it is complex and heterogeneous–varying over a wide range of spatial scales from nanometers to centimeters [1, 2]. It has been recently recognized that whole-brain imaging at an intermediate “mesoscopic” scale, consisting of functional neuronal units, could provide valuable new insights into normal and abnormal brain function [3, 4]. Magnetic resonance imaging (MRI) is uniquely suited for this purpose given its non-invasiveness and large field of view (FOV). Its nominal spatial resolution (≈ (2mm)^3^ voxels in clinical MRI scanners) however is poor by comparison to optical microscopic techniques with signals averaged over millions of cells and orders-of-magnitude more fine processes over an entire voxel. This limitation could be partly overcome using quantitative MRI methods which typically use a biophysical model to explain the MR signal arising from a voxel.

Diffusion MRI is one such method which analyzes the signal decay under a magnetic field gradient arising from diffusion of water molecules in complex media to infer its microstructure. The advent of strong, rapidly switching magnetic field gradients [5, 6, 7] has the potential to obtain whole brain MRI scans *in vivo* at near mesoscopic length scales in a clinically feasible time frame. However, there is also a need to develop accompanying models to relate the voxel-averaged measured macroscopic diffusion weighted (DW) signals to the underlying mesoscopic tissue structure and composition. Although many models have been proposed for this purpose [8, 9], the one with minimal assumptions while being rich enough to accurately describe whole brain mesoscopic tissue architecture is needed.

The diffusion tensor distribution (DTD) model [10] which assumes tissues to be composed of an ensemble of water compartments or pools each exhibiting anisotropic Gaussian diffusion, whose associated diffusion tensor ellipsoids may vary in size, shape, and orientation, as described by a probability density function (pdf) or a distribution of diffusion tensors within a voxel, *p*(D), (i.e., the DTD [11, 12]) is a possible candidate. These subcompartments, which can be viewed as microvoxels, are akin to the myriad of microscopic compartments observed in electron microscopy (EM) or optical images of brain tissue where water is assumed to undergo hindered diffusion. This assumption at the mesoscopic length and time scales accessible by the current MRI scanners is reasonable given the high water permeability of lipid plasma membranes (via passive diffusion across their lipid bilayers) [13, 14], the presence of water channels, such as aquaporins in astrocytes, and facilitated diffusion, which translocates water via the action of ion pumps [15, 16]. In addition, owing to the action of the glymphatic system in the brain, aqueous cerebrospinal fluid (CSF) is continuously being cycled rather than restricted within closed compartments to prevent build-up of toxins in brain parenchyma [17]. Experimental evidence also affirms Gaussian diffusion in brain parenchyma at mesoscopic length and time scales given that the mean apparent diffusion coefficient (mADC) reaches a constant asymptotic value at diffusion times longer than ∼10 ms [18, 19, 20] rather than continuously decreasing with diffusion time, characteristic of water being confined in pores or compartments (i.e., restricted). We have also recently shown that multiple-PFG signals, which are sensitive to mesoscopic heterogeneity [21] are independent of the diffusion time in live human brain, further strengthening the argument for assuming mesoscopic Gaussian diffusion [22].

Imaging the DTD non-parametrically (or empirically) therefore has the potential to reveal distinct mesoscopic water pools present within a voxel. However, inversion of the diffusion weighted MRI (DWI) signal data to estimate the non-parametric DTD is challenging for several reasons: 1) it requires taking a multi-dimensional inverse Laplace transform (ILT), which is well-known to be an ill-posed, ill-conditioned inverse problem, and 2) a highly under-determined one, as the number of unknowns in the high-dimensional space of diffusion tensors (i.e., 6D corresponding to the six independent components of the 2^*nd*^-order diffusion tensor) is several orders of magnitude larger than the number of DWI data points that can practically be acquired in a clinically feasible time period.

Several studies have attempted to estimate and map features of the DTD each having their attendant strengths and limitations. Westin et al. [23] used a cumulant expansion to express the MR signal in terms of our Gaussian DTD whose 2^*nd*^-order mean tensor and 4^*th*^-order covariance tensor [11] were estimated from the MR signal decay profile. This can however only be used over a narrow range of b-values and with specific experimental designs. Topgaard, et al. estimated a reduced 4D DTD in a phantom assuming cylindrical symmetry of the underlying microdiffusion tensors [24] and more recently, applied this symmetry constraint to measure the DTD in human subjects [25, 26]. To describe heterogeneity and microscopic anisotropy in the cerebral cortex, Avram et al., estimated 3D DTDs by assuming that the local cortical reference frame (i.e., coincident with cortical columns and layers) could replace the principal axes or directions of the microdiffusion tensors [27]. We previously estimated a constrained normal DTD which revealed cortical layers in excised brain tissue [28, 29, 30], although it is computationally intensive. Overall, most of these parametric methods can not capture the rich diversity of diffusion microenvironments in individual water pools throughout the brain. Recently, Song et al. reported a non-parametric method to estimate the full 6D-DTD, however using only single pulsed field gradient (sPFG) DWI data [31]. The use of only single-PFG data however renders the problem highly ill-posed since they can result in the same signal profile for a variety of microstructure motifs thereby limiting its applicability [23].

In another vein, a promising algorithm for performing non-parametric, multi-dimensional ILTs was introduced in a different context by Benjamini and Basser [32, 33], where the 1D marginal probability density functions (pdf) of the dependent scalar variables, *T*_1_, *T*_2_ relaxation times and mean diffusivity were estimated separately, and then used as constraints for the subsequent estimation/reconstruction of their 2D joint pdf using a technique known as marginal distribution constrained optimization (MADCO). This approach resulted in vastly reduced data acquisition requirements as well as increased precision and accuracy of the estimated 2D joint distribution when ground-truth spectra were provided for comparison. Due to a lesser number of unknowns and the high effective signal-to-noise ratio (SNR), the estimates of the marginal densities are more precise and accurate. This approach originally implemented for 2D diffusometry-relaxometry applications is increasingly more powerful as dimensionality increases given the non-linear increase in the volume of the parameter space of unknowns with respect to the number of dimensions. However, extending this approach to more than two dimensions and to a joint distribution of elements of a 2^*nd*^-order tensor as opposed to scalars (e.g., *T*_1_ and *T*_2_), such as what we encounter in DTD MRI, is not straightforward. In this latter case the off-diagonal components of the tensor are statistically coupled to the diagonal ones, making the estimate of their individual marginal densities or distributions directly from DWI data more challenging.

In this study, we present a new non-parametric or empirical approach that significantly extends the MADCO framework to estimate the entire 6D-DTD from a hierarchy of lower dimensional joint marginal densities of the elements or components of the diffusion tensor, which are estimated using DW signal data from a set of single, double, and triple-PFG MR experiments. The successive application of a hierarchy of marginal distribution of the DTD dramatically reduces the volume of the space of admissible solutions and reduces the computational time. We extensively test and vet this framework using simulated noisy DWI data generated using realistic DTD motifs that are expected in the brain to assess the fidelity of the estimated DTDs. We experimentally apply our approach in the human brain *in vivo*, and measure and map DTD-derived invariant quantities and their glyphs to visualize salient mesoscopic tissue features characterizing diffusion heterogeneity and microscopic anisotropy.

## 2 Materials and Methods

### 2.1 DTD estimation

The diffusion weighted MR signal, *S*(`B`), acquired with a diffusion encoding b-tensor, `B`, from an ensemble of non-exchanging diffusion tensors distributed according to the DTD, *p*(`D`), is given by [10],

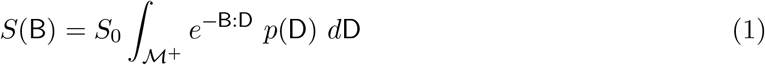

where *S*_0_ is the signal without diffusion weighting (i.e., *S*(`B = 0`)), `B:D` is the tensor dot product between the 2^*nd*^-order symmetric b- and diffusion tensors, respectively [34], and ℳ^+^ is the space of 3 × 3 positive semi-definite symmetric matrices. N.B. that the use of the kernel, *e*^−B:D^, explicitly assumes the subdomains within the voxel exhibit Gaussian anisotropic diffusion [28]. Alternatively, one can describe the equation above as a Fredholm Integral Equation of the First Kind in which one tries to solve for the distribution, *p*(`D`), from the signal, *S*(`B`). It can be observed that the tensor dot product of rank-1 b-tensors and rank-m (m > 1) diffusion tensors would result in redundant kernels making the inverse problem further ill-posed, hence single-PFG measurements alone are insufficient to estimate the full DTD.

The above signal model for a discrete set of diffusion tensors, `D`_*n*_, and experimental b-tensors, `B`_*m*_, is given by,

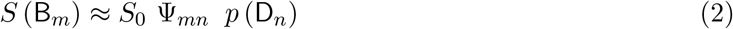

where Ψ_*mn*_ = exp (−`B`_*m*_ : `D`_*n*_) Δ`D`_*n*_ is the *m × n* anisotropic Gaussian diffusion kernel matrix, Δ`D`_*n*_ is the product of discretization step size of the independent components of the diffusion tensor, `D`_*n*_, and *p* (`D`_*n*_) is a *n*-dimensional vector consisting of the probability density of the discretized components of the DTD. Generally *n* ≫ *m*.

Since direct inversion is difficult, our approach to address this problem is to use a set of physically, mathematically, and statistically motivated constraints and *a priori* information to vastly reduce the space of admissible solutions for *p*(`D`). First, we use the positive definiteness constraint by zeroing diffusion tensors in the DTD that do not lie on the ℳ^+^-manifold as we have done in [28]. This results in approximately a 65% reduction in the degrees of freedom for the range of diffusivities used in this study (numerically estimated). Second, we use a hierarchy of lower dimensional (i.e., less than six) marginal distributions to constrain the 6D joint distribution estimation by partitioning ℳ^+^ into domains containing admissible solutions with higher probability while zeroing out others, thereby vastly limiting the remaining 35% of the solution space in ℳ^+^. The gradual pruning of the solution space by applying these constraints is graphically illustrated for two cases of 3D DTDs, for ease of visualization, involving two diagonal and one off-diagonal component in Figure 1, and all diagonal components in Figure 2.

**Figure 1:**
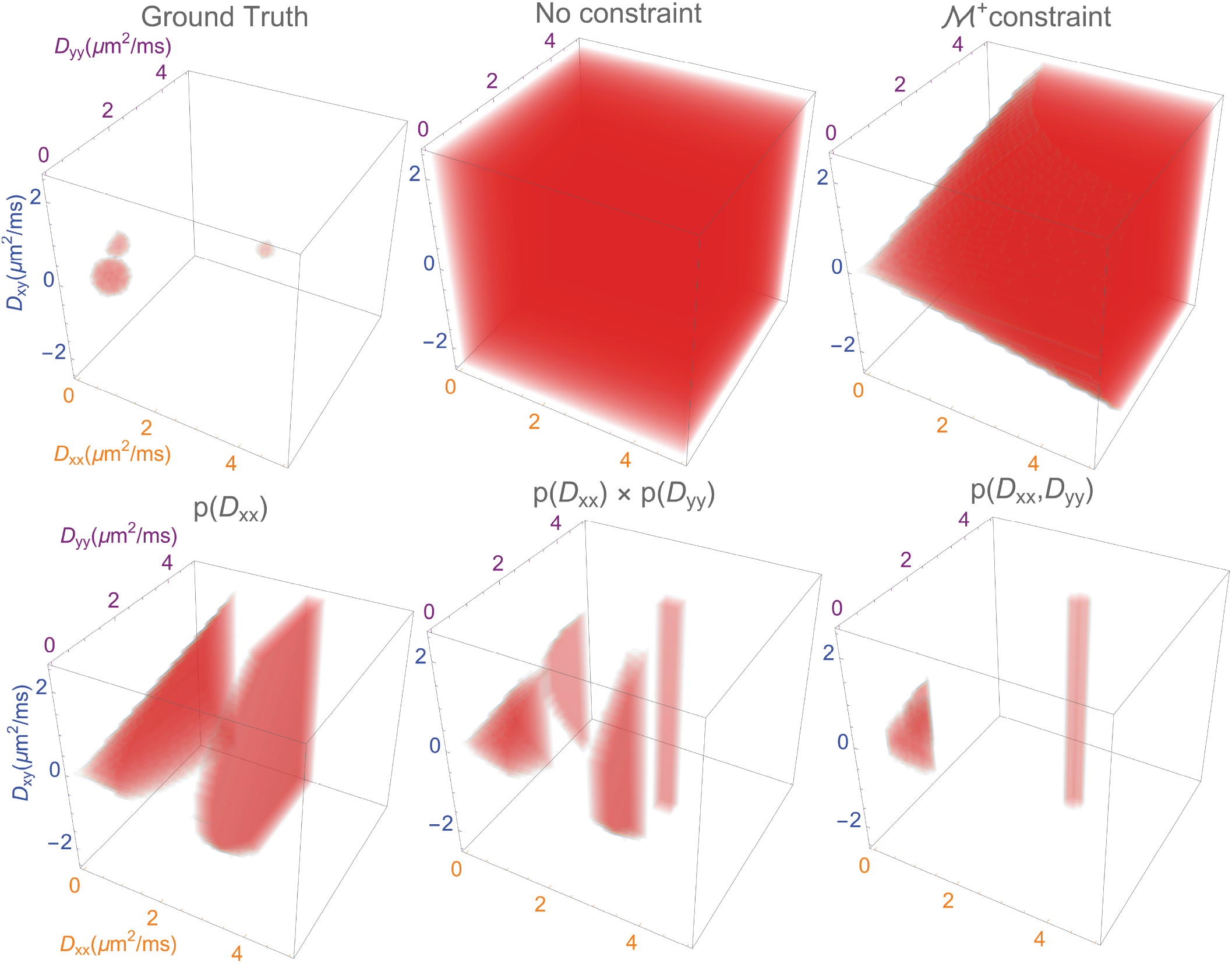
The gradual pruning of the solution space by the successive application of positive semi-definite (ℳ^+^) and marginal density constraints illustrated for the estimation of *p*(*D*_*xx*_, *D*_*yy*_, *D*_*xy*_). The domain of admissible solutions or region of interest (ROI) in the solution space (i.e., ℝ^3^) is shown in red. The ground truth DTD is assumed to be three probability spheres centered at *D*_*xx*_ = 0.2, 0.6, 3*µm*^2^*/ms, D*_*yy*_ = 1.4, 0.6, 3*µm*^2^*/ms, D*_*xy*_ = 0.2, 0.0, 0.0*µm*^2^*/ms* to represent white matter, gray matter and CSF respectively subject to the ℳ^+^ constraint. The vast 3D Cartesian solution space is first reduced by the application of the ℳ^+^ constraint as shown in the figure. A further reduction in the solution space is obtained by the application of 1D marginal density constraints for *D*_*xx*_ and *D*_*yy*_ and their 2D joint distribution.

**Figure 2:**
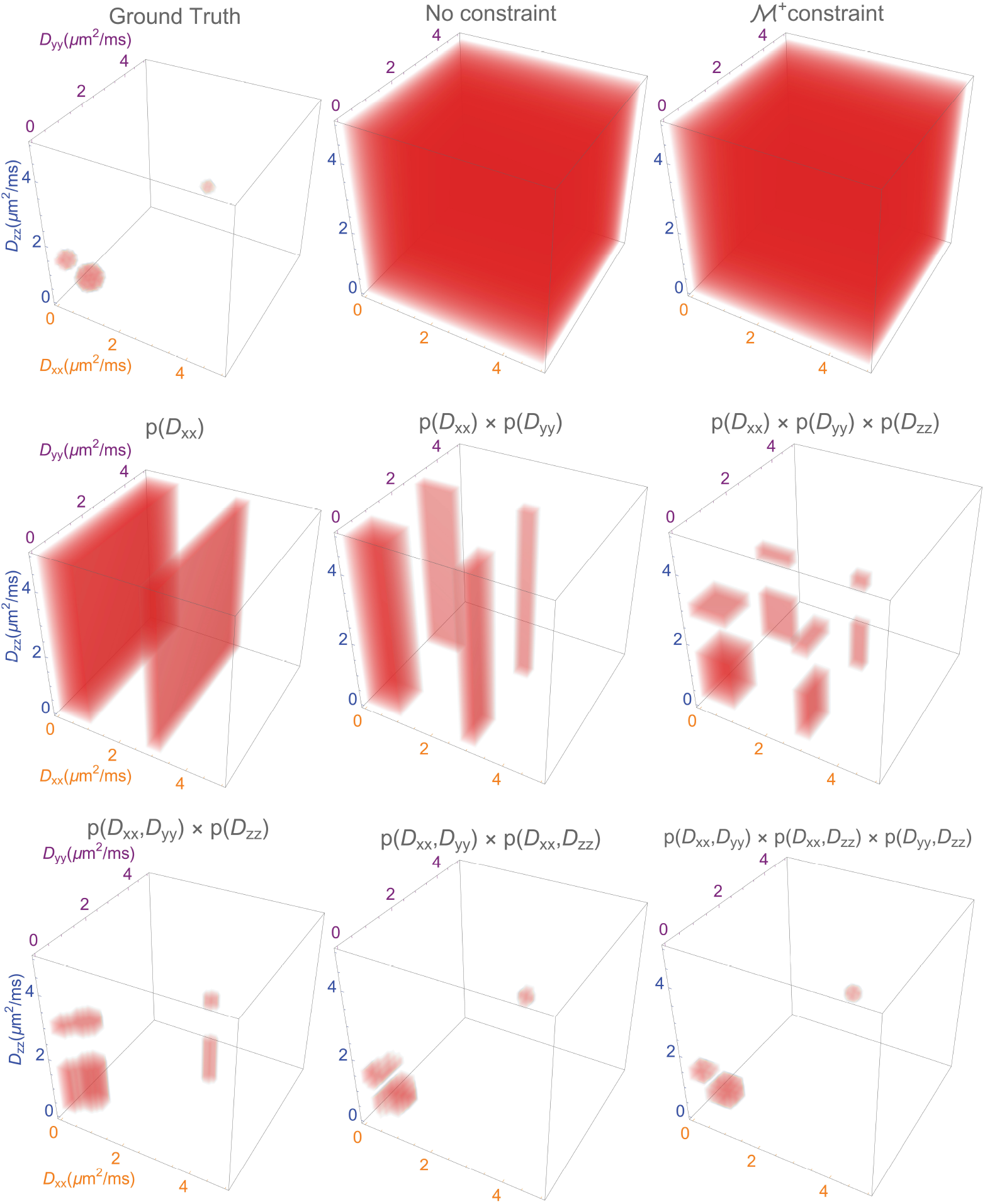
The successive pruning of the space of admissible solutions by the marginal density constraints is illustrated for the estimation of *p*(*D*_*xx*_, *D*_*yy*_, *D*_*zz*_). The regions of interest (ROIs) in the solution space (i.e., ℝ^3^) are shown in red. The ground truth DTD is assumed to be three spheres centered at *D*_*xx*_ = 0.2, 0.6, 3*µm*^2^*/ms, D*_*yy*_ = 0.2, 0.6, 3*µm*^2^*/ms, D*_*zz*_ = 1.4, 0.6, 3*µm*^2^*/ms* to represent white matter, gray matter, and CSF respectively. In this case, since all the diagonal elements of the diffusion tensor are by default set to be positive, the application of the ℳ^+^ constraint does not result in a volume reduction of the 3D Cartesian grid of unknowns. However, the application of 1D and 2D marginal density constraints result in a significant reduction in domain size as shown in the figure. The intersection of 1D marginal density constraints for *D*_*xx*_, *D*_*yy*_, *D*_*zz*_ shown in the second row results in a smaller sculpted domain in ℝ ^3^. This is further refined by applying the 2D marginal constraints, which results in a region very similar to the ground truth region, as shown in the last row.

Moreover, we recast this problem as one of convex optimization (Gasbarra, personal communication), guaranteeing unique solutions and exploiting efficient and robust algorithms to arrive at them. The resulting constrained convex optimization problem can be stated formally as follows,

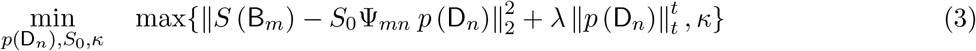

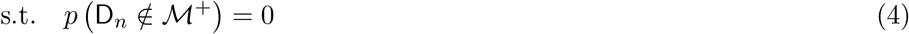

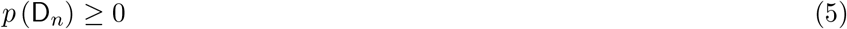

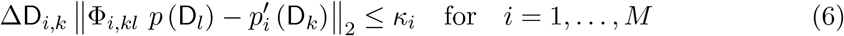

where *λ* is the Lagrange multiplier used to regularize the solution whose value is computed using the S-curve method [35], Φ_*i,kl*_ is the *k × l* matrix integral operator that maps the estimated DTD to its *i*^*th*^-marginal density of interest (N.B. the matrix multiplication involving Φ can become computationally and memory intensive operation if only 1D marginals are available as in the original MADCO implementation as *l* ≫ *k*), *M* is the number of marginals, *κ*_*i*_ is the tolerance variable which is introduced to relax the *i*^*th*^-marginal density constraint due to uncertainty in the estimation resulting from noise in the measurements, Δ`D`_*i,k*_ is the product of the discrete step size of diffusion tensor components for the *i*^*th*^-marginal density constraint, *t* is the norm used for regularization which was set to 1.5 to balance between *ℓ*_1_- and *ℓ*_2_-regularization [36], and *p*^*′*^ is the marginal density estimated independently as shown below. The pruning of the solution space (i.e., ℳ^+^) is accomplished by computing the cumulative density function (cdf) of the marginal densities and choosing those regions encompassing 95% of the distribution, which significantly reduces the rank of the Ψ and Φ matrices, making the problem computationally tractable. In words, Equations (3) to (6), we propose an estimation method of the DTD that is smooth and minimizes the error between the measured signal and the corresponding signal generated by the DTD model for that b-tensor, subject to the constraints that a) the diffusion tensors are positive definite, b) their densities are non-negative, and c) the computed marginal densities from the estimated DTD agree with their corresponding measured densities.

The full 6D DTD estimation is performed hierarchically as sixteen lower-dimensional estimation problems with many fewer unknowns. First, 1D marginal densities of the three diagonal components of the diffusion tensor are obtained from the 1D ILT via *ℓ*_*t*_-regularized non-negative least squares (NNLS) [35]. These were then used as constraints to successively reconstruct the 2D, 3D, 4D, and 5D marginal densities involving all observable combinations of diffusion tensor components (e.g., *p* (*D*_*xx*_, *D*_*yy*_), *p* (*D*_*xx*_, *D*_*yy*_, *D*_*xy*_), etc.) leading to the full 6D DTD. The lower dimensional projections of the 6D DTD are estimated from DW signals that selectively encode the diffusion tensor components of interest. The selectively encoded MR signal is inverted by solving Equation (3) to obtain the marginal density. The convex optimization problem for estimating the marginal and joint densities is performed using the CVXPY software package [37, 38] with the MOSEK solver [39].

The 6D DTD space was discretized into a Cartesian grid with the diagonal components of the diffusion tensor ranging conservatively from 0 to 6 *µm*^2^*/ms* to accommodate the range of mean apparent diffusion coefficients (mADCs) expected in live human brain, and to include the possibility for there being pseudo-diffusion in cerebrospinal fluid (CSF) leading to an mADC greater than that of free water at 37°C, 3 *µm*^2^*/ms*. The range of off-diagonal components was set to -3 to +3 *µm*^2^*/ms* since we do not expect values outside this range. The spectral resolution for all the components was set to 0.3 *µm*^2^*/ms*, which results in approximately 86 million unknowns needed to characterize the full 6D distribution without imposing any *a priori* information or constraints, making this problem otherwise computationally intractable.

### 2.2 Experimental design

We introduce a new experimental design, significantly extending our previous MADCO framework, which now hierarchically estimates all observable marginal densities and then uses them to estimate the full 6D DTD. While single-PFG measurements are straightforward to implement and are SNR efficient, they are not sufficient to measure correlations between and among different diffusion tensor elements since the generated rank-1 b-tensors do not fully span the space of b-tensors in ℳ^+^, nor are they amenable to estimating the covariance structure of the DTD [28]. For example, a diagonal b-tensor required to estimate the joint distribution of pairs of diagonal elements, (e.g., *p* (*D*_*xx*_, *D*_*yy*_)) cannot be realized with single-PFG data as it will inevitably also encode off-diagonal components. Thus, we use the single-PFG experiment to estimate the 1D marginal densities of only the diagonal components, double-PFG experiments to estimate the 2D/3D marginal densities involving pairs of diagonal components, and triple-PFG experiments to estimate the 3D/4D/5D distributions involving all the components of the diffusion tensor. Collectively, these constraints are used to estimate the 6D DTD. Previously, using our original implementation of MADCO [33], we employed only 1D marginal distributions of scalar MR parameters, such as *p*(*T*_1_), *p*(*T*_2_), and mean diffusivity *p*(*MD*), to estimate their 2D joint distribution. We have now extended MADCO to include a hierarchy of potentially all observable lower dimensional marginal distributions to further constrain the estimate of the highest dimensional (i.e., 6D) joint DTD, to make it accurate, feasible, and efficient.

The diffusion gradients in the single-PFG experiment are incremented individually such that the resulting b-values along a given axis are linearly spaced over a given interval. Selective encoding of the diffusion tensor components for the double-PFG experiment is achieved by zeroing the extraneous b-tensor components, which effectively reduces them to a 2 × 2 rank-2 matrix (or tensor) that is sampled randomly on ℳ^+^ using a compressed sensing approach. For the triple-PFG experiment, a set of 3 × 3 rank-3 b-matrices or tensors are chosen randomly on ℳ^+^ to observe the diffusion tensor components of interest while zeroing others, again using a compressed sensing approach.

A total of three sets of rank-1 (*n* = 11 for estimating the 1D marginal densities), six sets of rank-2 (*n* = 21, 31 for estimating the 2D and 3D marginal densities, respectively) and eight sets of rank-3 b-tensors (*n* = 31, 41, 51, 61 for estimating the 3D, 4D, 5D, and 6D marginal densities, respectively) with b-values ranging from 0 to 2 *ms/µm*^2^ were used to estimate the DTD. Thus a total of 11×3+21×3+31×4+41×3+51×3+61 = 557 b-tensors were used to reconstruct the full 6D DTD in each voxel with initially 86 million unknowns. The b-tensor rods/ellipses/ellipsoids used for hierarchically estimating the various marginal distributions are shown in Figure 3. It is clear that efficiently pruning the solution space is key to making the estimation of the DTD tractable.

**Figure 3:**
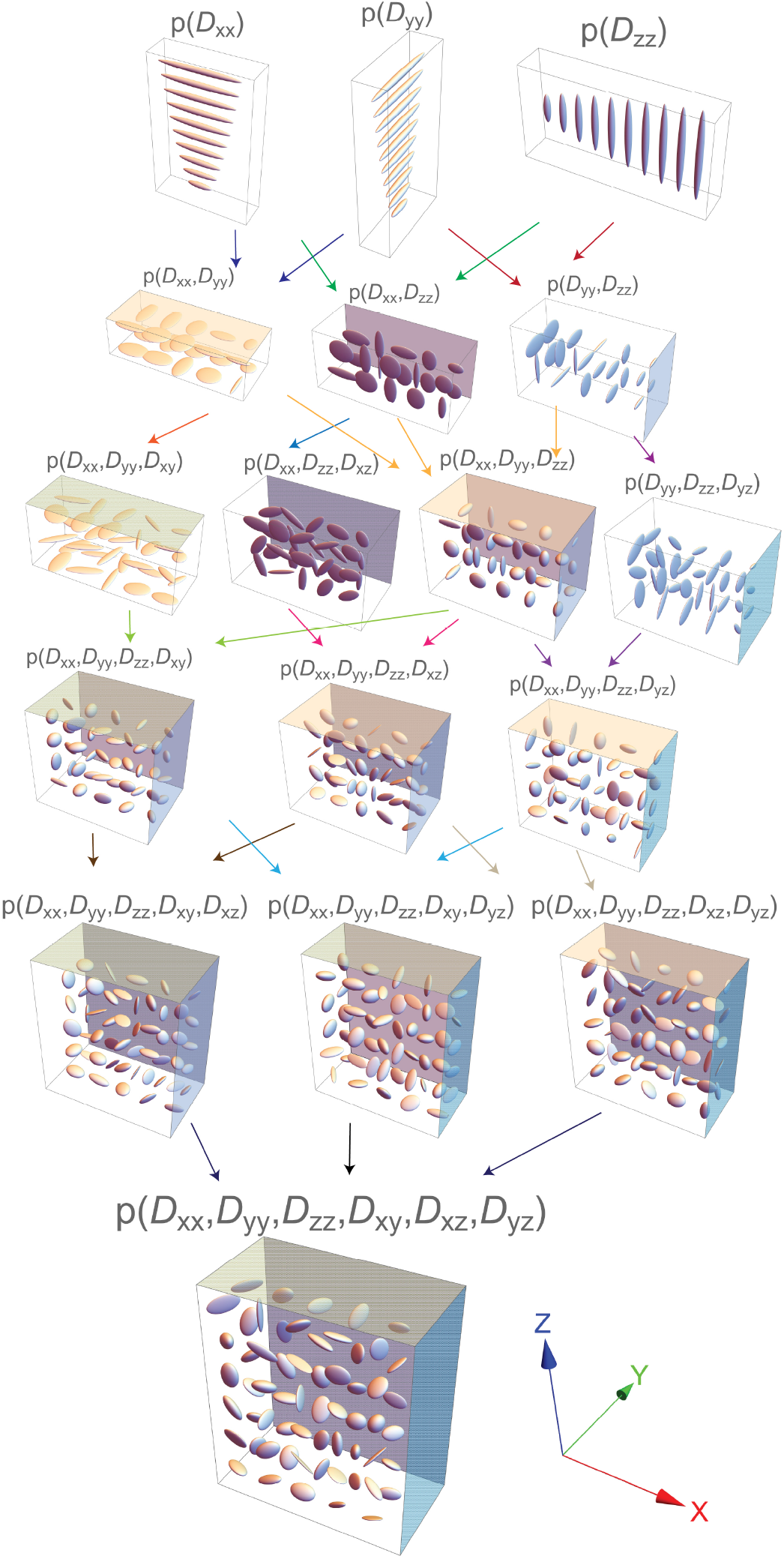
Hierarchical experimental design used to estimate the non-parametric 6D DTD depicted by plotting the ellipsoids of the sampled b-tensors for each set of DWI acquisitions. At the top are the rank-1 b-tensors displayed as sticks used to estimate the 1D marginal density of diagonal components of the diffusion tensor. In subsequent rows, the nD marginal distributions used to reconstruct the appropriate (n+1)D marginal distribution are shown using distinctly colored arrows ultimately leading to an estimate of the full 6D DTD. The estimated marginal distribution is indicated above each set of the b-tensors. To assist with 3D visualization, the plane where the b-tensors lie are indicated using shading of faces. In cases where the b-tensors are also rotated along the normal vector to the faces, a hatching pattern is used. For instance, the xy-plane is shaded in the estimation of *p*(*D*_*xx*_, *D*_*yy*_) while hatched in the estimation of *p*(*D*_*xx*_, *D*_*yy*_, *D*_*xy*_) to indicate the rotation around *z*-axis required to encode *D*_*xy*_.

### 2.3 Simulation of the DTD MRI pipeline

The efficacy of the reconstruction approach is investigated using a synthetic DTD phantom. The motif tested consists of equal proportions of gray matter, white matter and CSF. Gray matter is represented by an emulsion with spherical diffusion tensors whose MD is *γ*-distributed with a mean equal to 0.6 *µm*^2^*/ms* as that of brain tissue (i.e., 0.6 *µm*^2^*/ms* [40]). White matter is represented as two fiber populations whose ellipsoid orientations cross at 90° with MD = 0.6 *µm*^2^*/ms* and *D*_∥_ = 7.5*D*_⊥_ such as in coherent white matter fibers (e.g., corpus callosum) [40]. The CSF is represented using an emulsion of spherical diffusion tensors whose mean diffusivity is distributed according to a truncated normal distribution (i.e., only positive samples) with a mean close to that of free water at body temperature (i.e., 3 *µm*^2^*/ms*) [30]. The above motif was chosen since it is one of the most complex case expected in live brain tissue consisting of size, shape and orientation heterogeneity within the same voxel. The DW signal from this synthetic phantom was generated for the various b-tensors required with our experimental design using Equation (2). Gaussian noise was added to the real and imaginary channels to study the reconstruction fidelity on the effect of signal-to-noise ratio (SNR), which was varied from 150 - 600 for the non-diffusion weighted images.

### 2.4 MRI measurements and image pre-processing

MRI data were acquired using a 64-channel RF coil on a 3T clinical scanner (Prisma, Siemens Healthineers) with nominal gradient strengths up to 80 mT/m per channel and a 200 T/m/s slew rate using an echo planar imaging (EPI) readout. The measurement was performed on healthy volunteers (N = 3) who provided informed consent in accordance with a research protocol approved by the Institutional Review Board (IRB) of the Intramural Research Program (IRP) of the National Institute of Neurological Disorders and Stroke (NINDS).

DWIs were acquired with a field of view (FOV) = 210 mm × 210 mm × 120 mm, 2 mm isotropic spatial resolution at 1965 Hz/pixel bandwidth, and GRAPPA acceleration factor = 4. The b-tensors are realized using interfused-PFG (iPFG) MR pulse sequence that is capable of generating b-tensors of all ranks within a single spin-echo sequence, as shown in Figure 4 [30]. The sequence is versatile, SNR efficient, free from concomitant field induced signal loss, and possesses a shorter echo time (TE) as compared to traditional multiple-PFG sequences. The rank-1 b-tensors used in single-PFG measurements were acquired with *δ\*Δ = 15*\*32 ms resulting in a TR*\*TE = 6, 200*\*75 ms. The rank-2 b-tensors used in double-PFG measurements were acquired with *δ\*Δ = 14*\*40 ms resulting in a TR*\*TE = 6,000*\*85 ms. The rank-3 b-tensors used in triple-PFG measurements were acquired with *δ\*Δ = 15*\*57 ms resulting in a TR*\*TE = 8, 100*\*120 ms.

**Figure 4:**
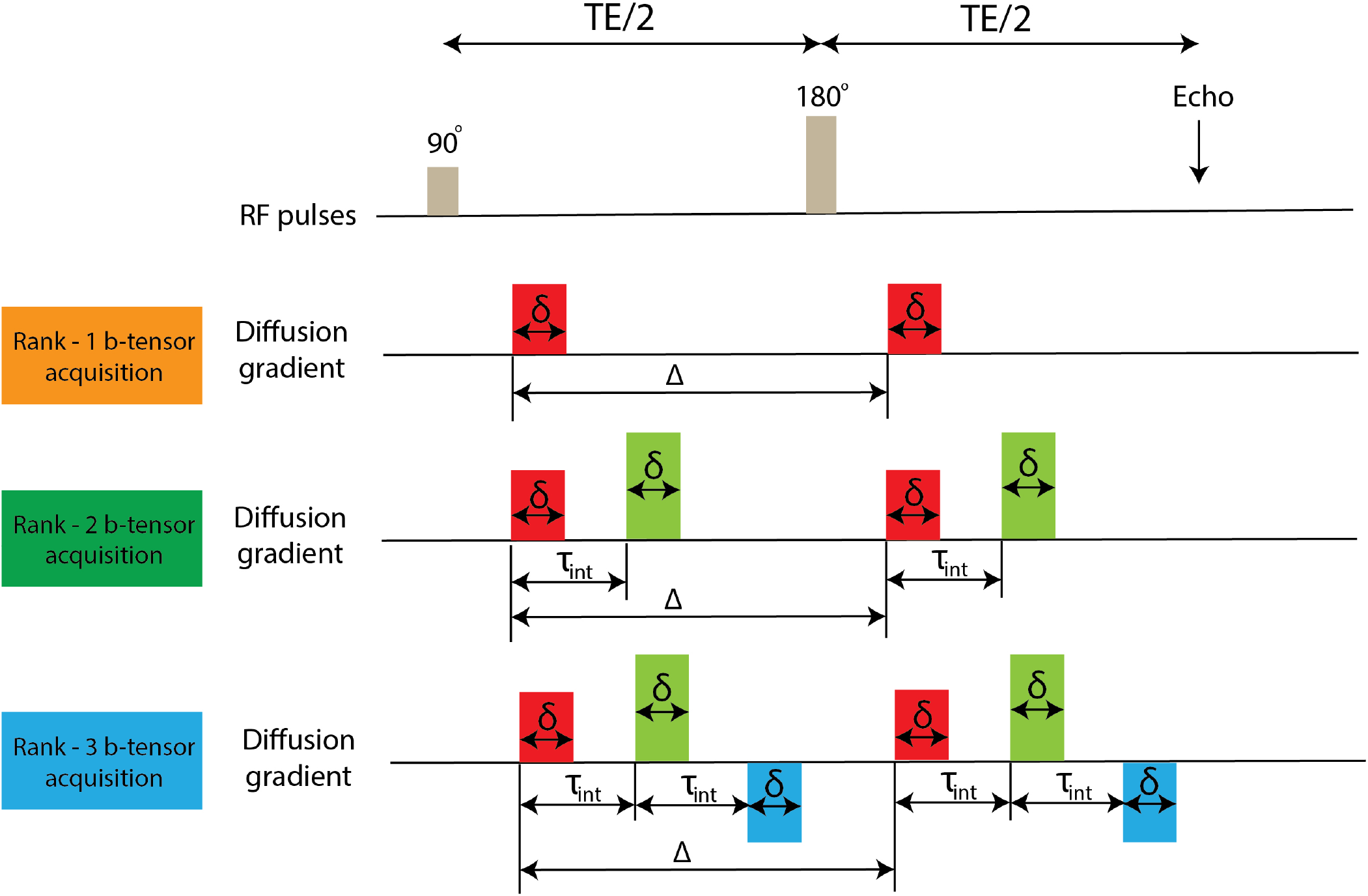
Multiple PFGs in a single DW spin-echo sequence used to estimate the non-parametric or empirical DTD. (First row) Single-PFG experiment used to measure the 1D marginal distributions of the diagonal components of the DTD. (Second row) Double-PFG experiment used to measure the 2D and 3D marginal distributions obtained from the DTD. (Third row) Triple-PFG experiment used to measure the 4D, 5D marginal distributions along with the full 6D joint DTD. The diffusion encoding gradients, *G*_1_, *G*_2_, *G*_3_, are shown in red, green, and blue, respectively. These pulsed-field gradient pairs may be applied along the same or different directions. The duration of the individual diffusion gradient pulses, *δ*, the diffusion time, Δ, and the interfusing time, *τ*_*int*_, are also indicated.

A *T*_2_-weighted fast spin-echo acquisition with 1 mm × 1 mm × 1.7 mm spatial resolution and TR*\*TE = 8, 000*\*72 ms was used to affine register the individual DWIs [41, 42] using the FSL software environment [43] to reduce the effect of sample motion thereby enabling cross data analysis. Gibbs ringing in the DWIs were reduced using the sub-voxel shift technique [44] implemented in the DIPY software environment [45]. The DWIs were also denoised using Marchenko-Pastur principal component analysis (MP-PCA) [46] implemented in MRTrix software environment [47] to reduce line broadening in *p*(D) by boosting the SNR. The resulting nominal SNR calculated using two *b* = 0 *s/mm*^2^ images [48] was approximately 160. Voxels with only CSF such as in ventricles were excluded using DTI due to large signal variations resulting from pulsations, flow, noise etc which would result in a dense spectrum. Tissue microstructure is further isolated by removing the free water compartment via truncating the *p*(D) whose MD is greater than 2.5 *µm*^2^*/ms* and re-normalizing it.

### 2.5 DTD visualization

To reduce the dimensionality of the DTD, it is prudent to study and visualize its distinct features using various scalar invariant measures, introduced for DTI [49], and informative glyphs, such as in [12, 28]. The size–shape distribution of the intravoxel diffusion tensor ellipsoids are described by the 3D iso-surface contours and their 2D projections of the distribution of its three orientation invariants, namely the norm (i.e., 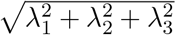 where *λ*_*i*_ is the *i*^th^ eigenvalue of the diffusion tensor), the fractional anisotropy (FA), and the mode of anisotropy (i.e., 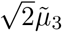 where 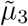 is the skewness of eigenvalues of the diffusion tensor) [50, 51]. The mode of anisotropy is weighted by the FA due to its noise sensitivity for spherical tensors. The orientation heterogeneity of the intravoxel diffusion tensors is described by the microscopic orientation distribution function (*µ*ODF) which is the ensemble-averaged ODF of micro-diffusion tensors within a voxel [30]. Similarly we also calculate the microscopic fractional anisotropy (*µ*FA) by computing the ensemble-averaged FA directly from the non-parametric DTD. Higher-order sample moments, such as the 2^*nd*^-order mean tensor, 4^*th*^-order covariance tensor, etc. are readily calculated directly from the empirical DTD as ensemble averages, as well. Of course, DTD MRI subsumes DTI so all DTI-derived parameters, such as the MD, FA, DTI-ODF, DTI-derived tracts, etc. can also be computed directly from the mean diffusion tensor, described in [28, 52].

The DTD is partitioned in each voxel using a non-parametric k-means clustering approach on the micro diffusion tensors. The number of clusters were empirically determined until degenerate clusters were observed. The above mentioned quantities can be computed for each distinct probability mode or distinct water pool to more accurately quantify mesoscale heterogeneity.

## 3 Results

The effect of SNR on the reconstructed DTD is investigated using a three-compartment synthetic phantom consisting of gray matter, white matter, and CSF in a single voxel as shown in Figure 5, which includes the ground truth results along with those obtained for SNR ranging from 150 - 600. The reconstructed DTD is visualized using 3D size-shape distribution isocontours as well as the computed 3D *µ*ODF. The first three moments (i.e., mean, variance and skewness) of the MD distribution along with FA and *µ*FA are plotted as a function of SNR. The ground truth DTD has three peaks in the size–shape distribution contours with the *µ*ODF showing the dominant crossing fiber orientations. The gray matter and CSF peaks were centered around 1 *µm*^2^*/ms* and 5 *µm*^2^*/ms* in size, respectively, and both had zero anisotropy indicating spherical diffusion tensors while the white matter peak was centered around 1.5 *µm*^2^*/ms* with anisotropy close to one indicating prolate diffusion tensors (N.B. size is given by the norm or determinant and not the Trace or MD of the diffusion tensor).

**Figure 5:**
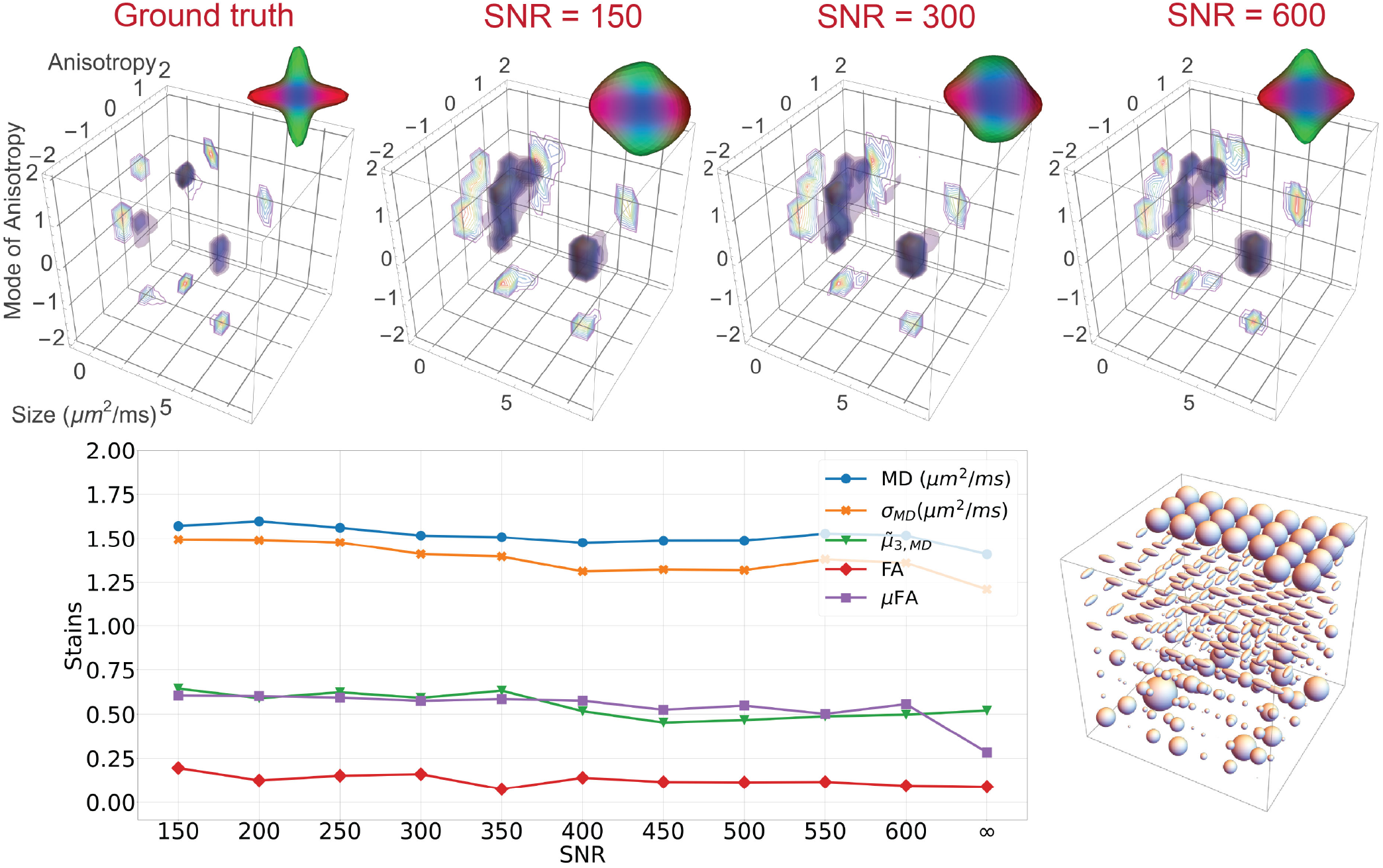
Effect of SNR on the reconstruction of a three-compartment motif within a single voxel composed of 1) white matter modeled as two sets of diffusion tensors whose ellipsoids cross at 90°, 2) cerebrospinal fluid (CSF) voxel modeled as an isotropic emulsion of diffusion tensors with MD larger than parenchyma, bracketing that of free water at body temperature, and 3) gray matter was modeled as an emulsion of spherical diffusion tensors with MDs smaller than in CSF. The motifs are displayed using micro-diffusion tensor ellipsoids. The 6D-DTD for each SNR is displayed using the size-shape distribution depicted using iso-surfaces or contours in the 3D parameter space of ellipsoid size, anisotropy, mode of anisotropy of the diffusion tensors, and the micro-orientation distribution function (*µ*ODF) computed from the DTD. The DTD-derived scalar measures characterizing the size (i.e., first three statistical moments of the MD distribution) and macro- and micro-anisotropy (i.e., FA and *µ*FA, respectively) are plotted as a function of SNR along with the ground truth values (i.e., SNR = ∞) to assess noise sensitivity. It can be observed that our approach is robust over a wide range of SNR values tested.

At SNR of 150, tissue and CSF peaks were separated but gray and white matter peaks within the tissue appear merged in the size-shape distribution plots. The *µ*ODF shows the two fiber orientations albeit smooth. The CSF peak is also shifted to a higher value than the ground truth. As the SNR increases, the CSF peak shifts closer to the ground truth while the gray and white matter peaks starts to separate. The *µ*ODF becomes sharper with increasing SNR while still showing the correct crossing fiber orientations, and approaching towards the ground truth. The stains were mostly stable over the range of SNR tested and closer to the ground truth value.

The various water pools along with their signal volume fractions derived from clustering the DTD obtained in three voxels from different parts of the brain tissue (corpus callosum, crossing fiber region and cerebral cortex) from a representative subject are shown in Figure 6. In corpus callosum, a single water pool was detected with the *µ*ODF pointing along the fiber orientation, and a broad size distribution was observed with fairly uniform FA and anisotropy mode close to one indicating prolate type tensors. In the crossing fiber region, four different water pools were detected. Three of these pools have similar size-shape distribution (prolate type tensors with size closer to tissue) but their *µ*ODF was pointing in orthogonal directions. The dominant water pool had smaller size but similar shape profile compared to the three other water pools and appeared to show two crossing fibers. The cerebral cortex had three different water pools. The dominant water pool has an one isotropic and slightly anisotropic diffusion tensors with size closer to that of the parenchyma. The *µ*ODF is inflated likely from the presence of the isotropic compartment but points normal to the cortical surface. The other two water pools were orthogonal white matter fibers of different sizes as shown by the shifts in the peak of the size distribution and their anisotropy mode.

**Figure 6:**
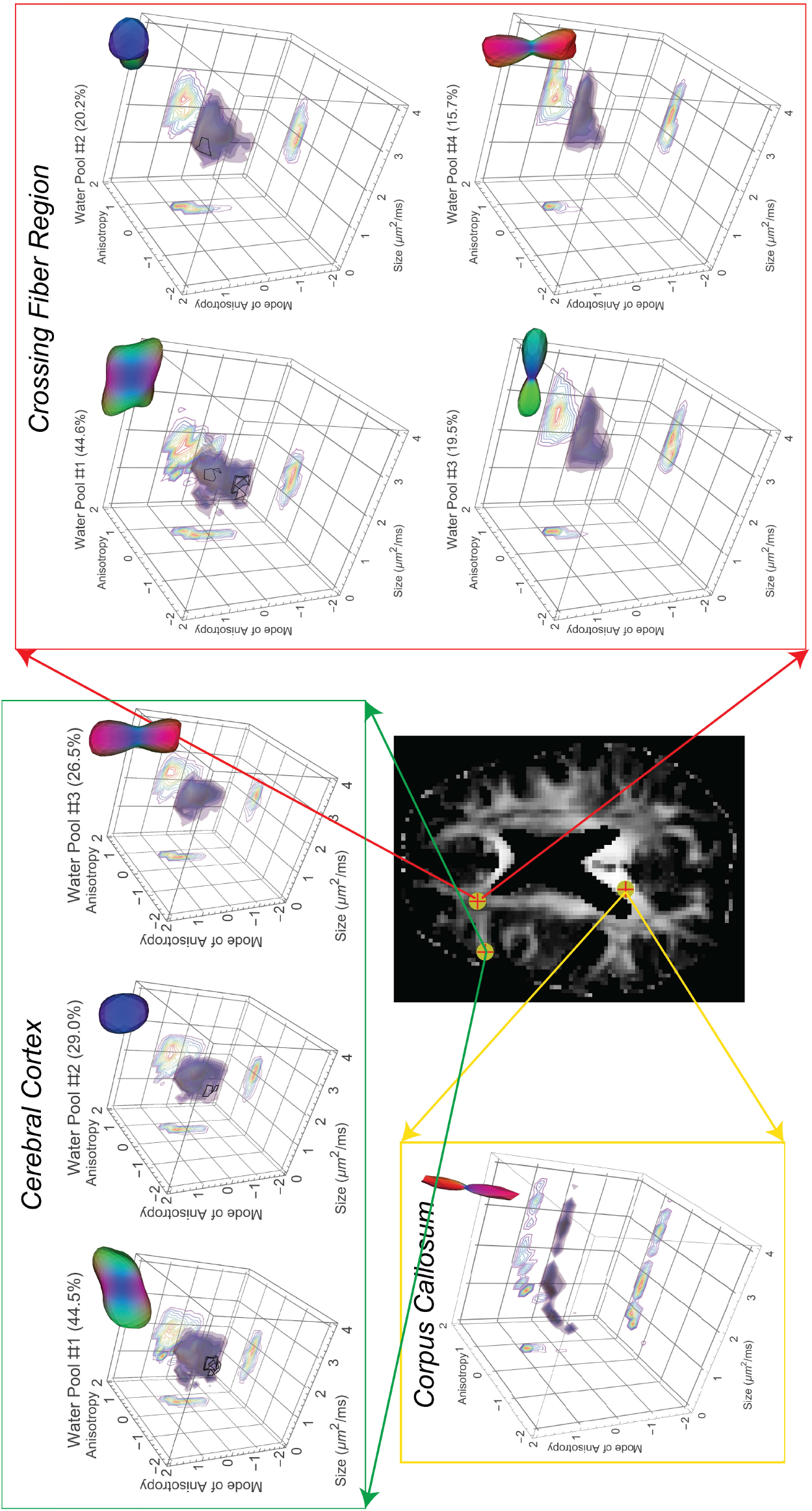
DTD reconstruction obtained in an axial human brain slice whose fractional anisotropy (FA) image is shown at the center. A set of three voxels were chosen in different brain regions (corpus callosum, cerebral cortex and crossing fiber region). The 6D-DTD is visualized using iso-surfaces or contours in the 3D parameter space of size, anisotropy, mode of anisotropy of the diffusion tensors, and by the micro-orientation distribution function (*µ*ODF) computed from ensemble averaging the DTD. The estimated DTD is clustered using k-means approach to reveal the diffusion properties of various water pools inside a voxel.

The various stains derived from the estimated DTD for an axial slice from a representative subject is shown in Figure 7. The maps include the non-diffusion weighted image, moments of ADC, and FA and *µ*FA maps. The MD map is fairly uniform as expected but the ADC standard deviation and skewness show new contrast. The ADC distribution in the corpus callosum exhibited high standard deviation but almost zero skewness while internal capsule fibers showed a positive skew with elevated standard deviations. The ADC skewness along the interface between tissue and CSF at the outer edge of the brain was predominantly negative. The contrast in macroscopic FA was as expected with corpus callosum having higher values and smaller values in crossing fiber regions. The microscopic FA recovered the lost FA in crossing fiber regions but showed new contrast in gray-white matter interfaces and in some white matter areas.

**Figure 7:**
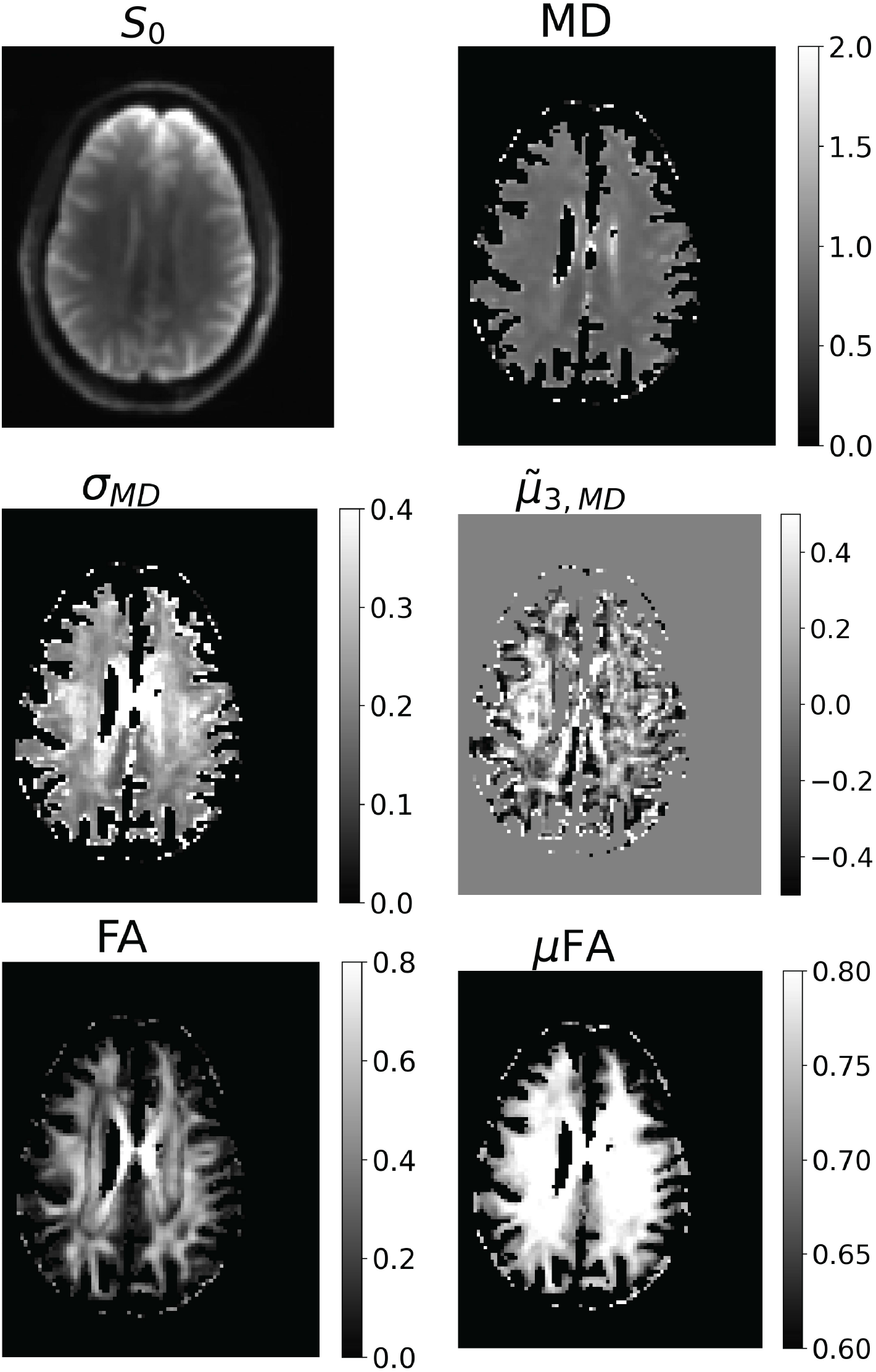
Maps derived from the estimated non-parametric DTD for an entire axial slice from a representative volunteer. This includes the non-diffusion weighted image (*S*_0_), moments of the apparent diffusion coefficient (ADC) such as its mean (MD), standard deviation (*σ*_*MD*_), and skewness 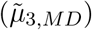, fractional anisotropy (FA), and microscopic fractional anisotropy (*µ*FA). The ADCs are expressed in *µm*^2^*/ms*. It can be observed that while MD is uniform, the standard deviation and skewness maps show new contrast among various white matter tracts. The *µ*FA shows new contrast and restores contributions of microscopic anisotropy that the FA powder averages.

The efficacy of free water elimination using our approach is shown for a single voxel in an axial slice of a representative volunteer containing both CSF and tissue compartments Figure 8. The large size peak corresponding to the free water is clearly removed in the size-shape distribution plot, and the extraneous peak in *µ*ODF is also removed which shows the free water pool is not necessarily isotropic. The resulting DTD more accurately reflects the tissue microstructure inside the voxel which contain a white matter fiber bundle.

**Figure 8:**
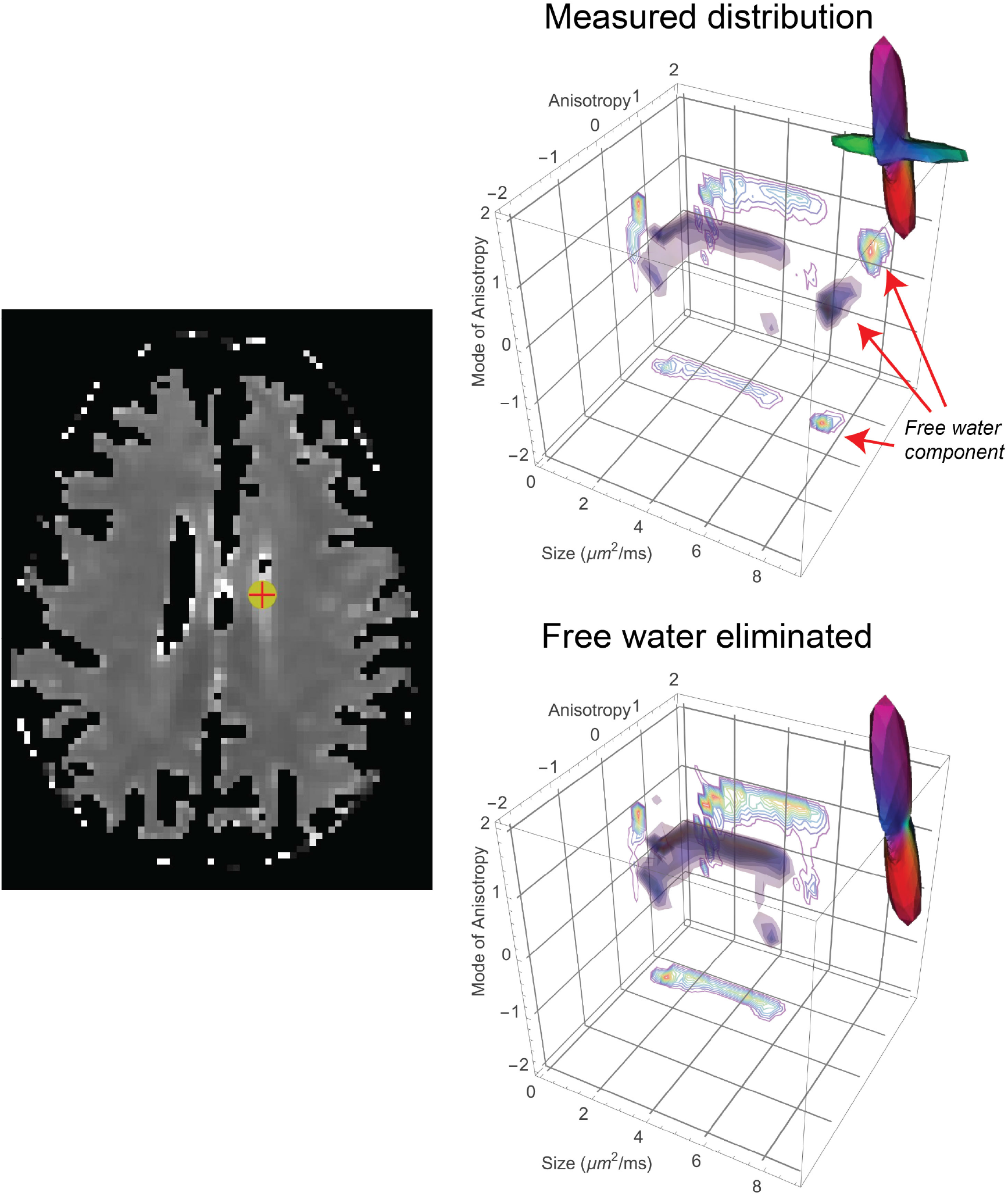
Free water elimination using DTD MRI for a voxel (indicated using a cross) in an axial human brain slice whose mean diffusivity map is shown at the left. Free water was eliminated by filtering the micro diffusion tensors derived from the estimated DTD whose mean diffusivity is greater than 2.5 *µm*^2^*/ms*. The 6D-DTD is visualized using iso-surfaces or contours in the 3D parameter space of size, anisotropy, mode of anisotropy of the diffusion tensors, and by the micro-orientation distribution function (*µ*ODF) computed from ensemble averaging the DTD. Removal of the free water component is visible in the size-shape distribution plot with the absence of the peak at large size shown using arrows. The extraneous peak in the *µ*ODF arising from CSF flow variation is removed reflecting the tissue microstructure that is partial volumed.

## 4 Discussion

In this study, we have demonstrated a MRI based methodology to measure and map mesoscopic water pools within live brain tissue. Diffusion in these water pools are statistically described using a non-parametric probability density function (pdf) of diffusion tensors (i.e., DTD) which is measured in each imaging voxel. The DTD is a six-dimensional pdf estimated using a hierarchy of marginal densities of different dimensions, which progressively shrink the space of admissible solutions. We have thoroughly vetted our framework using a set of realistic synthetic digital phantoms and applied our approach on live human brain tissue where it holds the promise of revealing mesoscopic brain architecture and detecting subtle changes in the brain which are often invisible in traditional MRI scans.

### 4.1 Validity of the DTD model *in vivo*

A key premise underlying the validity of this measurement approach is that over the range of experimental DWI parameters used (i.e., pulse gradient widths, diffusion times, diffusion gradient orientations and magnitudes), that the MR signal can be modeled as arising from a superposition of contributions from non-exchanging water pools, each described using an anisotropic Gaussian net displacement distribution so that Equation (1) holds. Specifically, the use of the anisotropic Gaussian diffusion (tensor) kernel, *e*^−B:D^, in the integrand of Equation (1) is assumed to be valid. While not the subject of this paper, we have demonstrated elsewhere [22] that at least over a clinical range of DWI parameters routinely used, and in particular, those used in this study, that this assumption does hold explicitly, justifying the form of the kernel in Equation (1). Briefly, the combination of tissue water homeostasis and permeability facilitated by plasma membranes, aquaporins, etc., makes it unlikely for cell and tissue water to be purely restricted at the length scales probed in this study [53], which would violate the use of the anisotropic Gaussian kernel for DTD estimation [22].

### 4.2 Effect of noise on DTD-derived parameters

Despite the constraints imposed by this new MADCO method, performing the ILT is fundamentally an ill-conditioned problem requiring *a priori* information to reduce the effects of noise in the data. In this study we have assumed the solution to be smooth as indicated by the *ℓ*_*p*_-norm regularization term in Equation (3) which numerically broadens the peaks in the 6D space akin to the line broadening effect in NMR spectroscopy. This affects the resolving power of distinct spectra as observed in the inversions of simulated data with low SNR.

It is important to note that noise affects some DTD-derived parameters differently from their DTI counterparts. As observed in the simulation, noise could shift and/or broaden peaks of the estimated DTD. Its effects are exacerbated on water pools with higher diffusivity such as in CSF because its signal decays quickly as a function of b-values resulting in more noise in the analyzed signal as a whole. This is the one of the reasons we masked out purely CSF voxels as their spectra are artificially broadened occupying huge swaths of the 6D space which makes it challenging to estimate at reasonable spectral resolution given our approach relies on the sparsity of *p*(`D`). Capturing purely isotropic voxels such as CSF or other liquids is also technically challenging in the presence of noise because the ensuing isotropic broadening of the spectra in 6D space will inevitably introduce artifactual anisotropic tensors.

Ways to mitigate these effects include obtaining real and imaginary parts of the signals and removing the Gaussian noise on each channel, using methods like Marchenko-Pastur (MP) PCA [46], or even increasing SNR by enlarging voxel size or using stronger gradients [5, 6, 7] to reduce TE or improving pulse sequence designs. We could also reduce the smoothing error by using advanced regularization methods such as elastic net [36] as opposed to *ℓ*_*p*_-regularization which we used for ease of implementation.

### 4.3 Curse of dimensionality

The advantage of multidimensional MR is the specificity it provides by separating features in a nD space, which are often indistinguishable in 1D. This however comes at the price of a non-linear increase in the number of degrees of freedom relative to the finite amount of data that can practically be acquired in a clinical setting–the so-called “curse of dimensionality”. Given that physical quantities are often sparse in nD space, our expanded MADCO framework provides enormous computational economy and an improvement in accuracy by resolving peaks at very high spectral resolution by pruning regions of admissible solutions based on the marginal densities. As additional constraints from a hierarchy of marginal distributions are applied, both the number of degrees of freedom and the volume of the solution space are markedly reduced so that admissible solution domain become confined and ever smaller, thus overcoming the curse of dimensionality. For instance, the number of unknowns needed to solve the simulation problem at the lowest SNR tested was reduced approximately by four orders of magnitude using our approach. It should be noted that we are not limited by the form of Equation (1). We can expand the number of variables by augmenting the kernel to include other relaxation processes, such as those governed by *T*_1_ and *T*_2_ decay. Then, the number of marginal distributions we need to compute increases rapidly with the number of dependent variables. We do not see using more marginal distributions/constraints as an obstacle with advent of high imaging throughput scanners [5, 6, 7]. Although more computations are required to simultaneously satisfy them, these added constraints further restrict the domain of admissible solutions, again helping to overcome the “curse of dimensionality”.

### 4.4 Water pools in the human brain

We have shown that our approach could decipher varying number of water pools present in each voxel. The number of water pools detected are however limited by the length scale of the microvoxels we are sensitive to which is determined by the size of the b-tensor (i.e., pore size 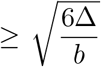), the width of the gradient pulse (i.e., pore size 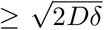), and the residence time of the spins within the microvoxel which should be less than Δ [30]. For the measurements presented in this work, the pore sizes we are sensitive are thus approximately 5 - 15*µ*m. At this resolution, we are sensitive to neuronal soma and axon fiber bundles but not individual axons.

The single water pool observed in corpus callosum with little shape heterogeneity but significant size heterogeneity is consistent with the known microstructure. The peak size was observed around 1.6 *µm*^2^*/ms* which corresponds to an ADC of 0.9 *µm*^2^*/ms*, for a gradient pulse duration used in this study, this corresponds to a pore size of roughly 5 *µm* which are present in this region albeit with low density [54]. In the crossing fiber region, our approach captured the three known orthogonal fiber populations known to be traversing this area (i.e., corpus callosum, corona radiata, and longitudinal fasiculus) [55] along with their relative signal fractions. The dominant water pool with slightly lower ADC than the other three water pools seems to suggest a mix of orthotropic and linear anisotropic tensors which may represent soma and dendritic processes. Cerebral cortex voxel was composed of both gray and white matter. The dominant water pool has *µ*ODF aligning with white matter bundle adjacent to it with orthotropic tensors likely arising from neuronal soma. There were also two orthogonal white matter water pools with different size and orientation distribution traversing this voxel.

### 4.5 Free water elimination

One confound that arises in our measurement of the diffusion properties of different tissue water pools arises from mobile “free water” pool. Several approaches have been used to try to eliminate this contribution to DWI signals, such as described in [56, 57, 58] so that parenchymal water diffusion can be analyzed separately from free water, which may exhibit unsteady flow, dispersion, or pseudo-diffusion, particularly in CSF compartments, e.g., in the ventricles. These methods often assume the free water has isotropic diffusion properties but it may exhibit artifactual varying anisotropic diffusion properties depending on their flow profile [59], which we have observed as well in voxels with a mix of CSF and tissue. Free water removal is a critical aspect of our implementation of DTD-MRI both in constructing the DTD and its DTD-derived parameters, so they represent parenchymal values. Here we first eliminate the free water signal contribution, which can be readily separated without invoking *ad hoc* criteria, retaining only the parenchymal water signal based on their mean diffusivity without making any assumptions of the anisotropy. The free water removal is facilitated by the high-dimensional character of the DTD data that relegates free water to well-defined, distinct domains of the 6D tensor space which makes our method more robust.

### 4.6 Future work

We are still exploring experimental design optimization strategies to improve the estimates of *p*(D), minimize the number of DWI acquisitions and total scan time, and increase accuracy (i.e., reduce statistical bias in estimated quantities), precision, and robustness. One objective is to determine the optimal number and mix of rank-1, rank-2, and rank-3 b-tensors for an optimal experimental design to facilitate radiological migration. An issue we plan to address and assess empirically is whether it is justified to assume the DTD comprises cylindrically symmetric “microtensors”, *a priori*, as is done in Topgaard et al. [24], resulting in 4D rather than 6D DTDs to estimate. Our pipeline now makes it possible to test this assumption.

A novel empirical or non-parametric DTD-MRI method was recently proposed by Song, et al. [31], which produced interesting and plausible *in vivo* MRI findings in live normal and cancerous human brain images acquired using the Connectome 1.0 MRI scanner. While encouraging, unlike our experimental design, they reported acquiring only single-PFG DWI MRI data, i.e., employing only rank-1 b-tensors. Compared to our highly constrained experimental design framework, this 1-D data would at most allow estimation of only three individual marginal distributions, *p*(*D*_*xx*_), *p*(*D*_*yy*_), and *p*(*D*_*zz*_), as depicted by the top row of Figure 3, but no other individual or joint marginal distributions. We plan to investigate the robustness, precision, and accuracy of estimates of the DTD when using only low-dimensional single-PFG data, but our assessment is that such data would only explore solutions on the outer boundaries of ℳ^+^ and not sample the interior of the domain.

Given adequate SNR, the benefit of estimating *p*(D) empirically, particularly in voxels containing multiple tissue types, (e.g., white and gray matter, CSF, and/or crossing white matter pathways) having distinct water pools, is that we expect to find multimodal DTDs whose distinct probability clouds are centered about different mean diffusion tensors (i.e., points) within the 6D tensor space whose statistical variability we can probe individually. One way to glean and summarize sub-voxel microstructural information in this case is to fit parametric distributions to these distinct probability clouds in 6D tensor space. The decomposition of the DTD into a family of parametric distributions can be used to summarize features of distinct intravoxel motifs in a principled manner.

## 5 Conclusion

In this work, we have presented a principled empirical framework to address the ill-posed, ill-conditioned inverse problem of estimating the DTD in each voxel within an imaging volume, significantly by extending our prior MADCO acquisition and analysis pipeline to employ a hierarchy of mathematical, statistical, and physical constraints and *a priori* knowledge that greatly limit the domain of admissible solutions. Here we also employ a novel experimental design, using single, double, and triple pulsed-field gradient (PFG) DWI acquisitions. We vet this framework with a digital diffusion MRI phantom to simulate DWI data possessing brain-like diffusion properties before applying it to *in vivo* human brain DWI data. Maps of DTD-derived intrinsic invariant measures and glyphs that depict salient mesoscopic tissue characteristics, revealing new microstructural features and details not reported previously in *in vivo* MRI studies.

## 6 Acknowledgments

We would like to thank Dario Gasbarra and Alexandru V. Avram for thoughtful discussions. This research was supported [in part] by the Intramural Research Program (IRP) of the *Eunice Kennedy Shriver* National Institute of Child Health and Human Development (grant number 1-ZIAHD008971-07), and National Institute of Neurological Disorders and Stroke, NIH. The contributions of the NIH author(s) are considered Works of the United States Government. The findings and conclusions presented in this paper are those of the author(s) and do not necessarily reflect the views of the NIH or the U.S. Department of Health and Human Services. This work was also partly funded by the Military Traumatic Brain Injury Initiative (MTBI^2^) in the Department of War (DoW) under award, HU0001-22-2-0058. This work utilized computational resources of the NIH HPC Biowulf cluster (http://hpc.nih.gov). The authors have no conflicts of interest to disclose. The views, information or content, and conclusions presented do not necessarily represent the official position or policy of, nor should any official endorsement be inferred on the part of, the Uniformed Services University, the Department of War, the U.S. Government, or The Henry M. Jackson Foundation for the Advancement of Military Medicine, Inc.

## Author Contributions

KNM and PJB conceived of, designed the research plan and drafted the manuscript. KNM wrote the MRI pulse sequences, and performed the measurements and DTD analysis. JES helped perform the MRI acquisitions. All authors carefully reviewed and edited the manuscript.

